# Is it time to retire Δ?

**DOI:** 10.1101/2025.06.13.659563

**Authors:** Roman A. Zubarev

## Abstract

The paradigm postulating that tissue stable isotopic ratios (*δ*_tissue_) equal those of the diet (*δ*_diet_) plus a small, quasi-constant isotope discrimination factor Δ emerged in the late 1970s, establishing stable isotope analysis as a dietary proxy. Today, this framework is still widely used across multiple branches of science, despite growing contradicting evidence. Here, we reanalysed several well-controlled laboratory experiments and universally found that Δ not only varies strongly with *δ*_diet_, but also changes sign at a certain *δ*_eq_ = *δ*_diet_ = *δ*_tissue_ value, which we term the Δ-equilibrium. The Δ-equilibrium phenomenon results from the sub-unity slope of the linear regression between *δ*_diet_ and *δ*_tissue_ and leads to converging of *δ*_tissue_ values. The most frequently observed position of the Δ-equilibrium on the (*δ*^13^C, *δ*^15^N) plane is (−21±1‰, 12±1‰). These findings firmly establish that stable isotopes are not neutral spectators but active participants in biochemical processes. If presented evidence holds in a much broader study, the paradigm *δ*_tissue_ = *δ*_food_ + Δ can finally be retired after half a century of service, being replaced by *δ*_tissue_ = *a*×*δ*_food_ + Const, where *a* is the newly defined isotope assimilation factor. The Δ-equilibrium position is then found as *δ*_eq_ = Const/(1 - a). The reason for isotope convergence remains a subject for future research, but likely hypotheses include evolutionary adaptation and isotopic resonance.

## Introduction

There are many reasons in research to seek objective information on diet, including studies on animal ecology [Hobson KA, 2023] and archaeology [Roberts P, 2002]. Climate studies also often rely on historical records inferred from animal diets [Fernández-García M et al., 2024]. In human health, there is a pressing need for more objective dietary information, as strong empirical diet-disease associations observed in controlled laboratory settings may be compromised when dietary assessments rely on self-reporting, which is not always reliable [O’Connell TC et al., 2012].

After the discovery of stable isotopes in the 1930s and until the late 1970s, it was generally believed that the isotopic composition of tissues reflected that of the diet, following the paradigm “you are what you eat”:

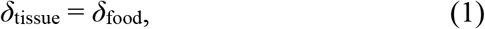

Here, *δ*represents the relative deviation, expressed in per mil (‰), of stable isotope ratios (SIRs) from a standard. The SIRs of carbon, hydrogen, nitrogen, oxygen, and sulfur vary naturally in the foods we eat and can be measured in a variety of tissues, as well as in blood, feces, urine, and exhaled air. Thus, according to paradigm (1), it was sufficient to analyze SIRs in any body tissue or excretion to infer dietary information.

However, after the seminal work of DeNiro and Epstein [DeNiro BN & Epstein S, 1978], this paradigm shifted. They provided empirical evidence that isotopic ratios in tissues are systematically different from those in the diet. They termed this difference the isotope discrimination factor (Δ) and postulated a new paradigm:

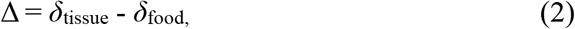

where Δ is a quasi-constant, resulting from the metabolic processes involved in element assimilation. DeNiro and Epstein concluded that animals are, on average, enriched in ^13^C by about Δ = 1‰ relative to their diet, while individual variations of Δ among animals raised on the same diet can differ by ±2‰. Knowing this, DeNiro and Epstein postulated that it is possible to perform dietary analysis based on δ^13^C determination in animals. They later extended this approach to ^15^N measurements [DeNiro BN and Epstein S, 1981].

The isotope discrimination factor Δ remains a cornerstone of modern dietary determination approaches. The prevailing belief underlying both paradigms (1) and (2) is that heavy isotopes, at least at abundances near natural levels, are neutral observers in metabolic assimilation processes. Consequently, isotope discrimination is presumed to be independent of the diet’s isotopic composition, and variations in Δ should be statistically unrelated to those of *δ*_food_.

However, Caut et al. (2009) analyzed hundreds of published datasets and found, for most organisms, a statistically significant negative correlation between these two values. They proposed the Diet-Dependent Discrimination Factor (DDDF) method, in which Δ is calculated from *δ*_food_ through linear regression. Although Caut et al. did not directly challenge paradigm (2), their results undermined its basic assumptions.

This study and the DDDF approach have been met with considerable criticism. For example, Auerswald et al. (2010) argued that Δ, being defined as dependent on *δ*_food_, will always negatively correlate with it even in the absence of a functional relationship between the two variables. Caut et al. counter-argued, and the debate remains unsettled.

We verified the conclusion of Auerswald et al. and found it valid in principle. However, the statistical link turned out to be quite specific. Although understanding this specificity allows, in principle, the use of linear regression to detect a hypothetical functional relationship between Δ and *δ*_food_, the presence of residual negative correlation makes any such effort potentially open to criticism. Therefore, here we chose a different test of paradigm (2).

According to both (1) and (2), linear regression between *δ*_tissue_ and *δ*_food_ should have a slope of 1, while the intercept should be equal to zero in (1) and Δ in (2). That the condition

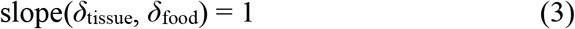

is essential for the paradigm (2) is illustrated on **Figure 1A**, where Monte Carlo simulations are shown for the case of slope of 0.5 and an intercept of -10‰.

**Figure 1.**
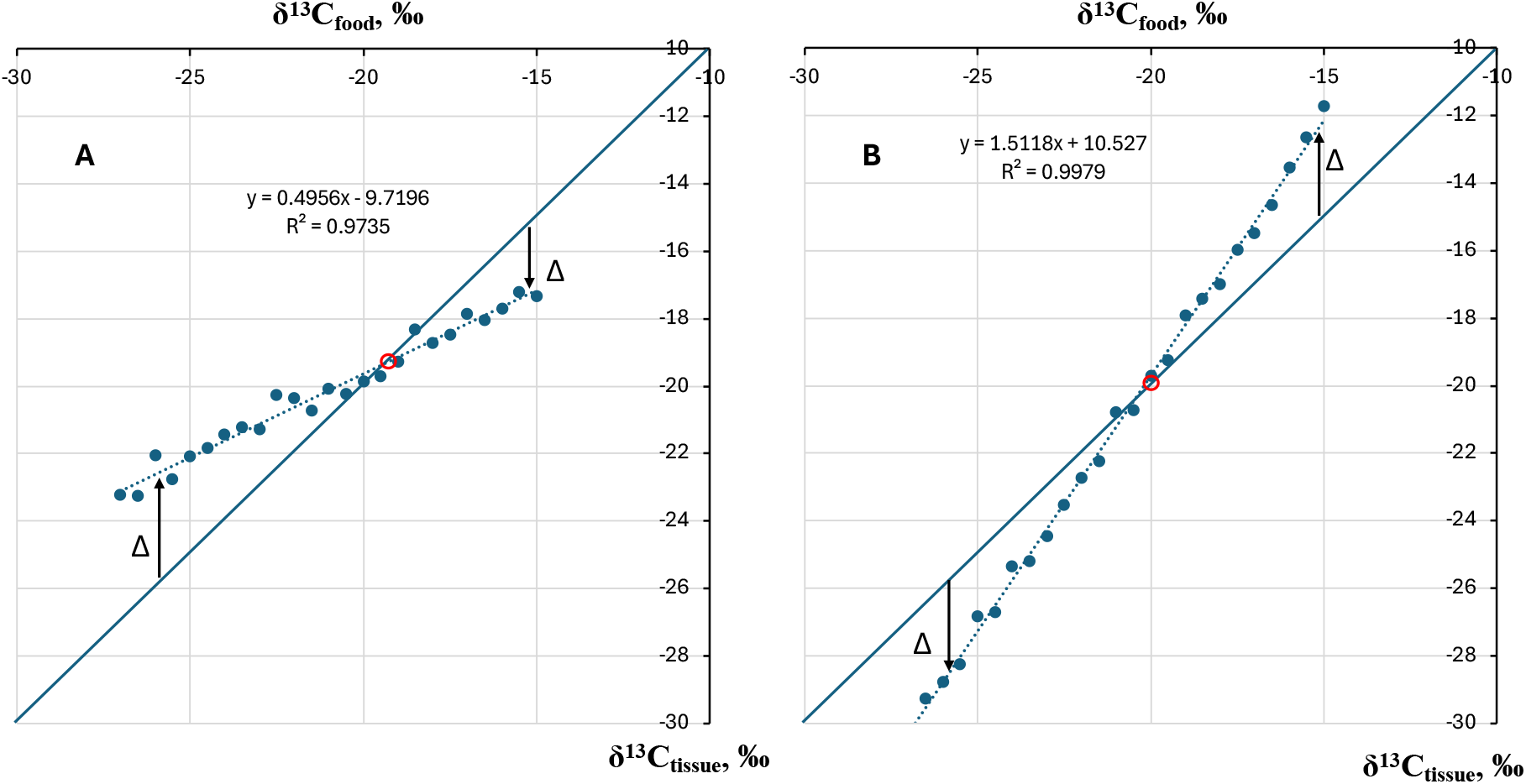
Monte Carlo simulations of a linear regression of the *δ*^13^C_tissue_ dependence from *δ*^13^C_food_ with: **A** - slope 0.5 and intercept -10 ‰; **B** - slope 1.5 and intercept 10 ‰. The diagonal line corresponds to *δ*^13^C_tissue_ = *δ*^13^C_food_, as in (1). Note how Δ magnitude and sign changes with *δ*_food_; the point of change is marked by red circle.

On the plot, the discrimination factor (Δ) varies with *δ*_food_ in both magnitude and sign, directly contradicting the assumption of a quasi-constant Δ. To the left of the transition point (marked by a red circle in **Figure 1A**), tissue is enriched in heavy carbon, resulting in a positive Δ. On the right side, however, heavy carbon is depleted, and Δ becomes negative. At the equilibrium point where Δ changes sign — referred to as the Δ-equilibrium — there is no isotope discrimination.

When the regression slope between *δ*_tissue_ and *δ*_food_ exceeds unity, as illustrated in **Figure 1B**, the equilibrium point persists, but the regions of heavy isotope enrichment and depletion are reversed. The coordinates of the Δ-equilibrium (*δ*_eq_) are identical on both axes and can be determined by the intersection of the lines *y* = *x* and y = *a*·*x* + *b*:

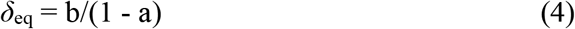

Three theoretical scenarios arise from this framework. First, if the regression slope is exactly unity, as predicted by paradigms (1) and (2), then Δ = *b*. Second, if the slope is less than unity, a Δ-equilibrium exists as defined above. In this case, the range of *δ*_tissue_ values is narrower than that of *δ*_food_, indicating that variations in dietary SIRs are partially buffered in the tissue, likely due to physiological mechanisms. The stability of the Δ-equilibrium can be quantified as 1−*a*, reaching its maximum when the slope is zero, at which point *δ*_tissue_ remains constant regardless of *δ*_food_.

The third scenario arises when the regression slope between *δ*_tissue_ and *δ*_food_ exceeds unity. In this case, the range of *δ*_tissue_ values becomes broader than that of *δ*_food_. Here, the Δ- equilibrium point is inherently unstable, as some mechanism amplifies dietary SIR changes in tissues by a factor equal to the slope. This amplification suggests that tissues not reflect but exaggerate dietary isotope variations, showing greater variability in tissue isotope values.

These three scenarios differ fundamentally in how isotopes participate in biochemical processes. Only in the first scenario, where the slope is unity, do isotopes act as passive tracers, reflecting dietary inputs without influencing their own discrimination. In contrast, in the second and third scenarios, isotopes modulate their own discrimination through underlying physiological or biochemical mechanisms. This modulation leads to either stabilization (slope < 1) or destabilization (slope > 1) of the Δ-equilibrium, indicating that isotopes are active participants rather than neutral observers in these processes.

To assess which scenario predominates in practice, we reviewed several well-cited datasets from the past two decades. Three datasets are based on controlled laboratory experiments with rodents and fish, each involving at least four distinct diets. An additional dataset comes from a field study on migrating birds, included to validate laboratory findings under natural conditions. Although these studies covered a variety of tissues, our analysis focused on muscle and liver — organs with high metabolic rates and rapid equilibration with dietary SIRs. We systematically extracted carbon and nitrogen isotope data for both *δ*_diet_ and *δ*_tissue_ from the published texts, tables, and, when necessary, figures, and conducted a comprehensive analysis to determine the prevailing scenario

## RESULTS

In the relatively recent work by Kadye et al. [Kadye WT et al., 2020], wild-caught Mozambique Tilapia *O. mossambicus* individuals were split randomly in 5 groups, with each group fed for up to 120 days an isotopically different diet. The expectations based on paradigm (2) and literature data were that for all diets and tissues discrimination factors will be Δ^13^C = 1.0 ± 2.0‰ and Δ^15^N = 3.4 ± 2.0‰ (**Figure 2A**). In reality, all isotopic data for muscle were confined to a stunningly compact cluster (**Figure 2B**).

**Figure 2.**
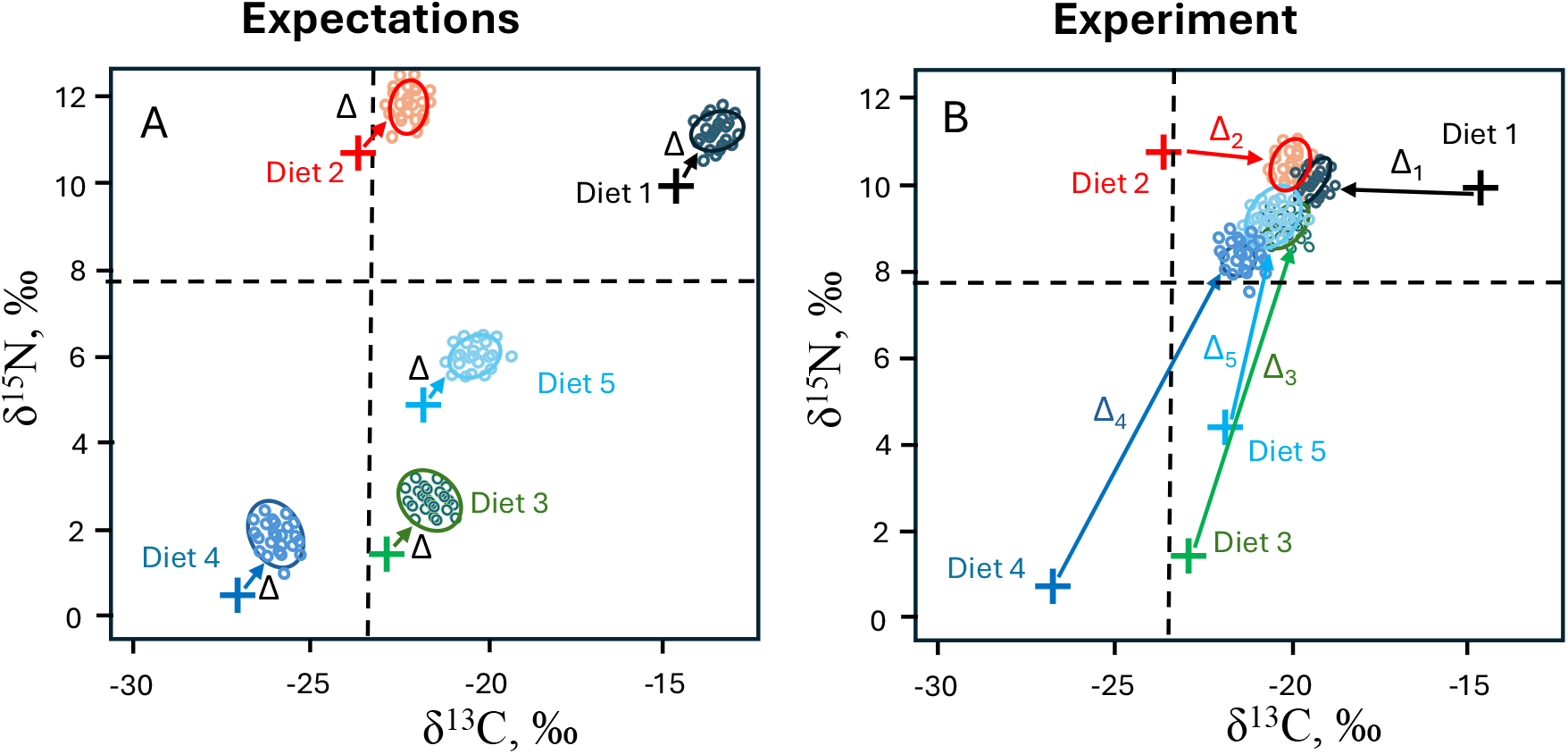
Isotopic compositions of diets and muscle of *O. mossambicus*. **A** – expectations based (2) with similar diet-tissue discrimination factors Δ for all experimental diets and tissues. **B** – actual measurements for muscle. Adopted from [Kadye WT et al., 2020].

In linear regression, the slope was only 0.16, in gross violation of (3b), while R^2^=0.86 certified that great majority of ^13^Cδ_tissue_ variations was accounted for by ^13^Cδ_diet_. The Δ- equilibrium point was reached for carbon at -19.9‰. For ^15^N, the slope was also very low, 0.20, while R^2^ was even higher, 0.94; Δ-equilibrium was reached at 10.8‰.

To assess the statistical significance of our findings, we conducted Monte Carlo simulations using the observed diet range and randomly assigned Δ values with magnitudes up to 10‰ for each diet. The simulations revealed that the probability of observing a tissue data cluster as compact as that shown in **Figure 2B** is less than 0.00003. This extremely low probability indicates that such a tight clustering of tissue isotope values is highly unlikely to result from random, independent variations in isotopic fractionation across different diets. Instead, these results strongly suggest the presence of a systemic mechanism causing isotopically diverse diets to produce isotopically similar tissue values.

This interpretation is further supported by earlier influential work, such as a study involving 48 male Norwegian rats, each six weeks old, which were fed for eight weeks on seven isotopically distinct but energetically equivalent diets (A–C, D:A+B, E:A+C, F:B+C, and G:A+B+C) [Caut S & Angulo E et al., 2008a]. At the conclusion of the experiment, the rats were euthanized and their tissues analyzed for isotopic composition.

While the spread of the tissue SIR data on **Figure 3** is larger than that in **Figure 3B**, the sizes of polygons created by the tissues are smaller than those of the diets, confirming the effect of isotopic conversion in tissues. The slopes of the δ^13^C_tissue_ vs δ^13^C_food_ linear regression for muscle and liver are 0.59 and 0.64, respectively, while high R^2^ values (0.88 and 0.99, respectively) testify to the strong diet-tissue link, especially for liver. Similarly, ^15^N slopes are 0.70 and 0.72, with very high R^2^ values of 0.97 and 0.98, respectively. From (4), the positions of Δ-equilibria on the (δ^13^C, δ^15^N) plane are (−22.1‰, 13.4‰) for muscle and (−20.8‰, 15.8‰) for liver.

**Figure 3.**
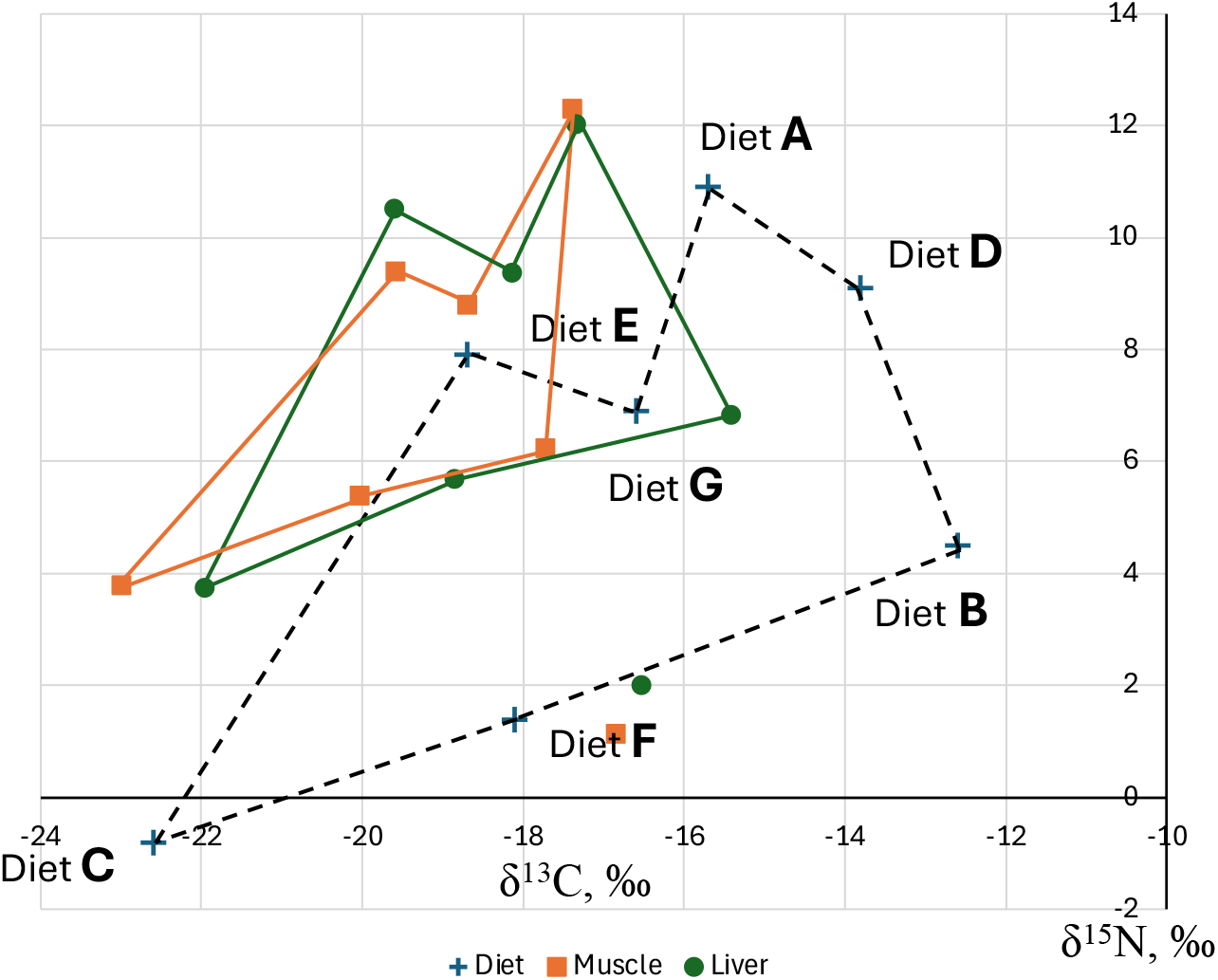
Isotopic compositions of the diets A-G, as well as muscle and liver of the rats grown for 8 weeks on these diets. Adopted from [Caut S et al., 2008a].

In yet another study, Lv et al. fed groups of Sprague–Dawley rats (n=19 in each group) four different diets for 21 days [Lv W, Ju T et al., 2012].

The results (**Figure 4**) demonstrate an extreme case of isotope convergence in tissues for carbon, with the slope of only 0.05 for muscle and 0.005 for liver, while the respective R^2^ values are 0.55 and 0.0008. At such small slopes, formula (4) produces large errors and the average value of -19.6‰ and -19.2‰ are better estimates for the position of Δ-equilibrium for muscle and liver, respectively. For ^15^N, the regression slopes are 0.24 for muscle and 0.52 for liver, with respective R^2^ of 0.99 and 0.97. According to (2), the δ^15^N coordinates of Δ- equilibrium are 6.8‰ and 13.7‰, respectively.

**Figure 4.**
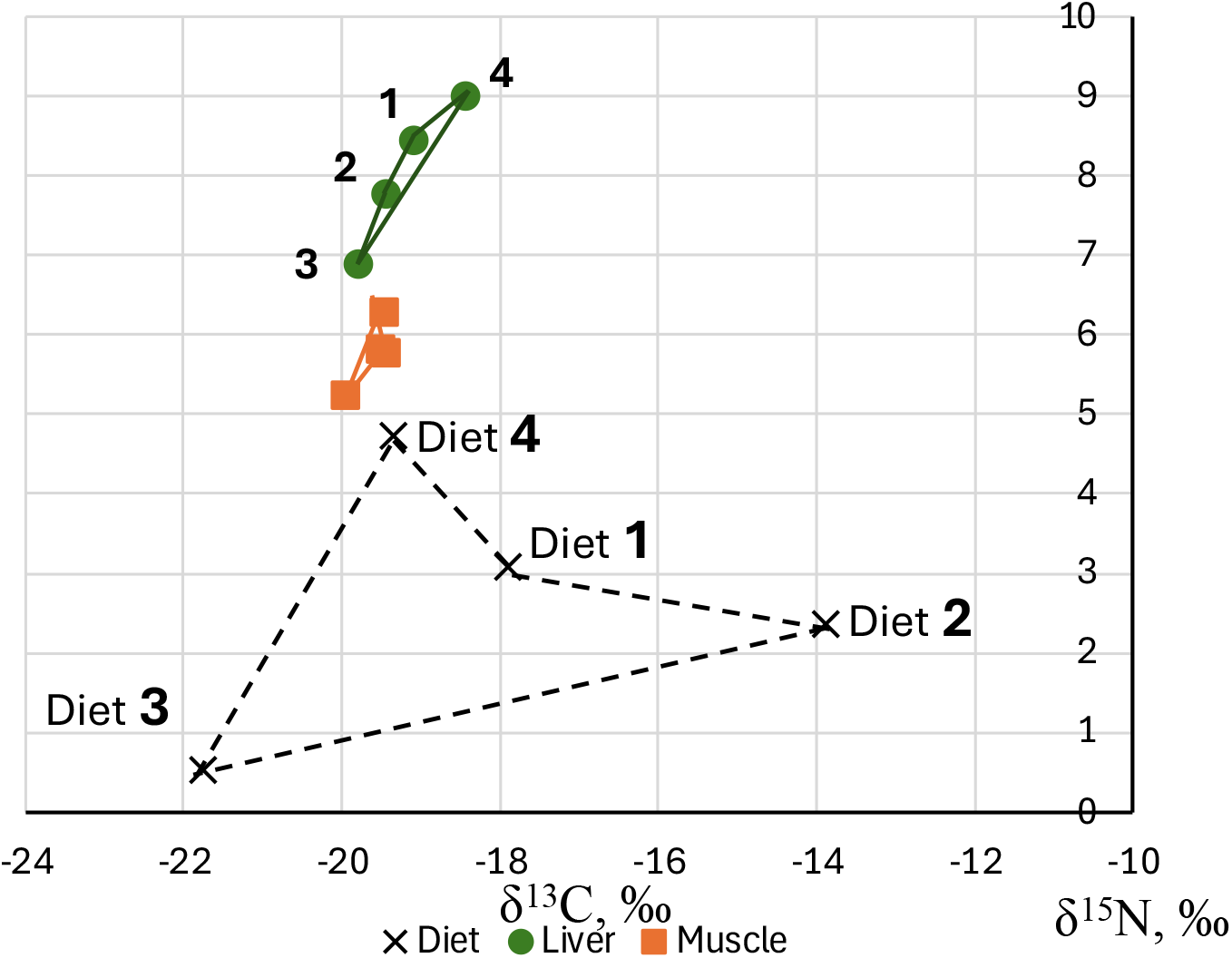
Isotopic compositions of the diets 1-4, as well as muscle and liver of the rats grown for 21 days on these diets. Adopted from [Lv W, Ju T et al., 2012].

Across all three laboratory studies reviewed, the data consistently contradict paradigm (2) and instead support the second scenario, where the slope of the linear regression between *δ*_tissue_ and *δ*_food_ is less than one. This means that SIRs in tissues tend to converge toward the Δ-equilibrium, showing less variability than the SIRs in the diet. Such convergence indicates a buffering mechanism in tissue isotope composition, rather than a direct, proportional reflection of dietary isotope values.

To address criticisms that Δ regression studies rely heavily on laboratory data, these findings were tested using a field dataset involving 132 individuals from 16 Chilean bird species. In this study, carbon and nitrogen SIRs were measured in muscle, liver, and crop contents (used as a proxy for “diet”) [Sabat P et al., 2013]. The regression slopes of *δ*^13^C_tissue_ vs. *δ*^13^C_food_ were 0.32 for muscle and 0.47 for liver, with R^2^ values of 0.57 and 0.38, respectively. For *δ*^15^N, the slopes were 0.41 (muscle) and 0.48 (liver), with R^2^ values of 0.55 and 0.62. These below-unity slopes, observed alongside moderate-to-strong R^2^ values, confirm that even in wild populations, tissue SIRs are less variable than dietary SIRs and converge toward a Δ-equilibrium.

The coordinates of the Δ-equilibria in the (*δ*^13^C, *δ*^15^N) plane were found to be (−22.1‰,8.6‰) for muscle and (−20.4‰,11.3‰) for liver, closely matching the laboratory results.

Motivated by these similarities, the positions of the Δ-equilibria for muscle and liver from all four studies were mapped onto a single plot. The resulting cluster was remarkably compact (**Figure 5**), suggesting a universal physiological mechanism governing tissue isotope composition. Although the exact statistical significance is difficult to quantify, even if the possible space for Δ-equilibria were as large as the entire plot, the probability of observing such a tight cluster by chance is less than 0.001. This strongly supports the conclusion that tissue SIR convergence is a systemic phenomenon, not a random occurrence.

**Figure 5.**
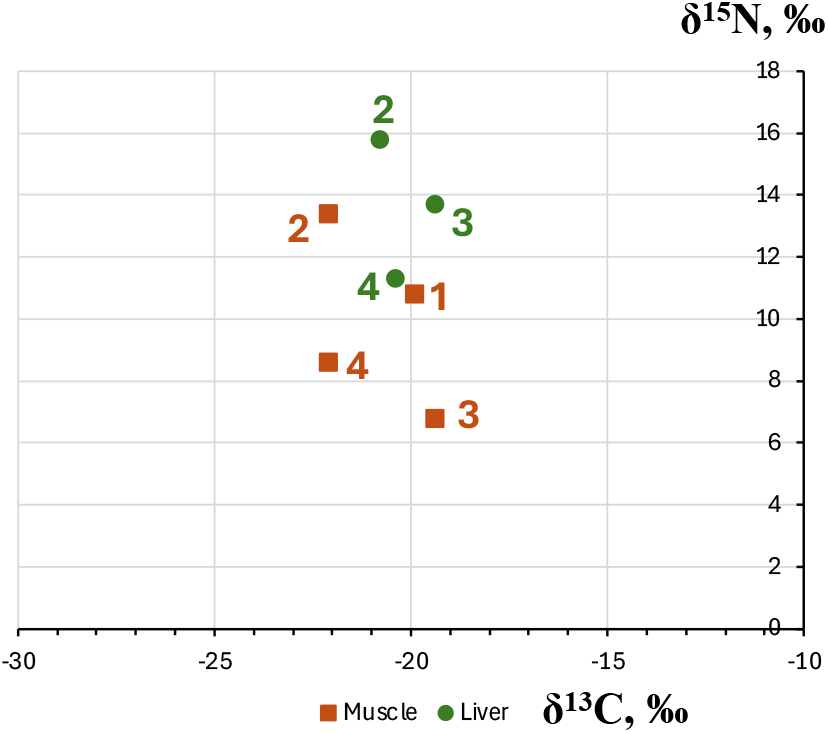
The positions of Δ-equilibria for muscle and liver in: 1- [Kadye WT et al., 2020], 2 - [Caut S et al., 2008a], 3 - [Lv W et al., 2012], 4 - [Sabat P et al., 2013].

Given the absence of statistically significant differences between muscle and liver data for both carbon and nitrogen, these data can be considered part of the same statistical group. Consequently, all 58 muscle and liver data points from the four studies were pooled for a combined linear regression analysis (**Figure 6**). The resulting models exhibited high R^2^ values and very low p-values (p < 0.0001 for carbon and p < 0.001 for nitrogen), confirming the statistical robustness of the regression for this dataset. The fitted regression lines intersect the *δ*^13^C_tissue_= *δ*^13^C_food_ line at *δ*^13^C = −20.5‰ and *δ*^15^N=11.4‰, which define the Δ-equilibrium positions for these tissues.

**Figure 6.**
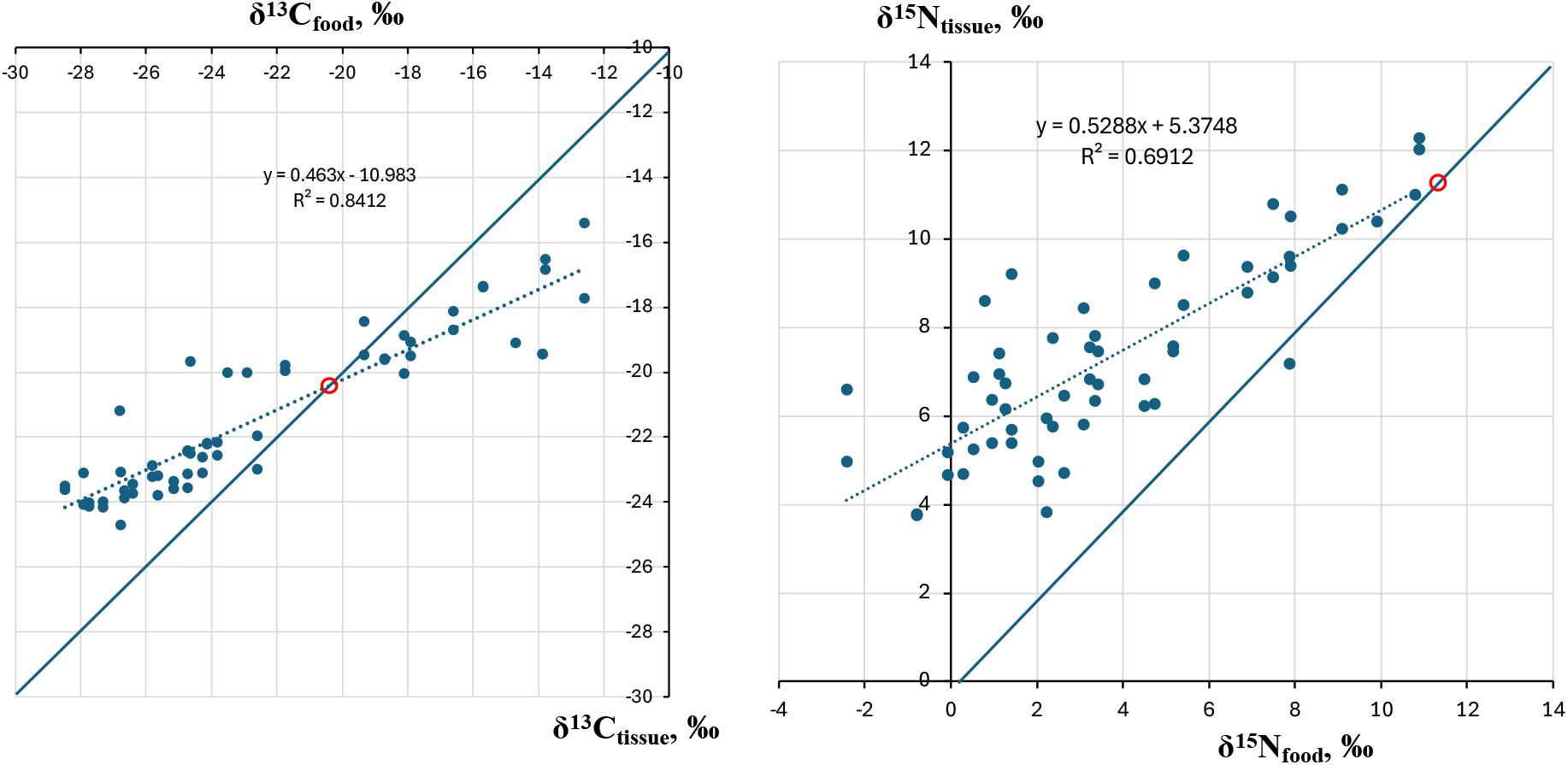
Combined plot of 58 muscle and liver datapoints from the four studies in **Figure 5**. The diagonal line corresponds to δ^13^C_tissue_ = δ^13^C_food_, as in (1). The equilibrium points (δ^13^C = -20.5‰, δ^15^N = 11.4‰) are marked by red circle.

Converting these data to Δ vs δ_food_, obtain the plots shown in **Figure 7**. The regressions, despite possibly being affected by the residual correlation [Auerswald K et al., 2010], cross zero at the positions of Δ-equilibrium in **Figure 6**, which largely validates the approach by Caut et al. Not surprisingly, their regression for carbon in mammals crosses zero at a similar value, -18.9‰ [Caut S, Angulo E et al., 2009]. The regression for nitrogen, being less statistically significant, crosses zero at a higher value than ours.

**Figure 7.**
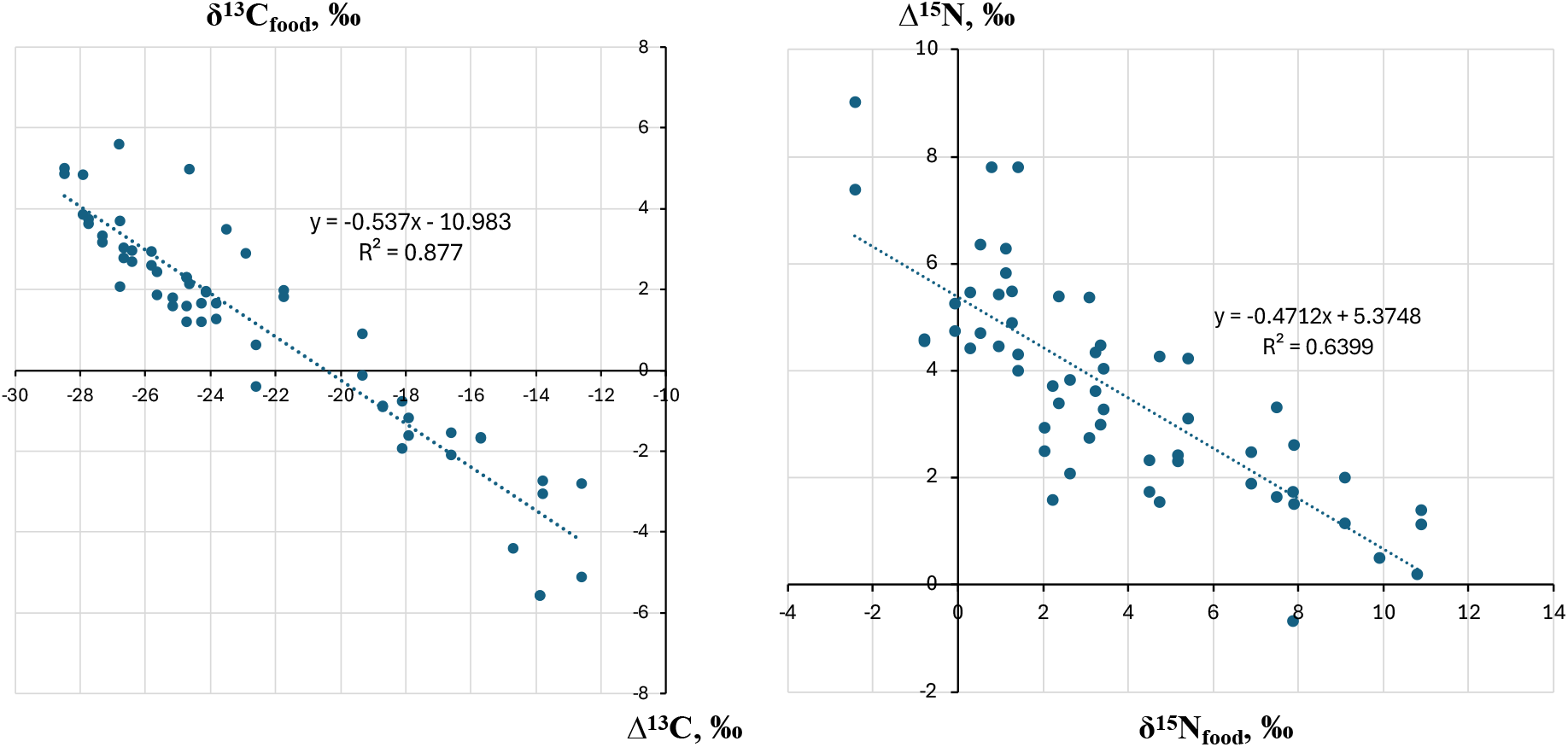
Linear regressions of Δ versus δ_food_, data from Figure 6.

To further investigate the residual correlation between Δ and *δ*_food_ as reported by Auerswald et al., we conducted analogous Monte Carlo simulations. These simulations consistently produced a slope near -1 and an R^2^ value around 0.5, as predicted by paradigm (2), where variability in Δ is equally explained by *δ*_diet_ and *δ*_tissue_. However, in our empirical data (**Figure 7**), the observed R^2^ values were notably higher than 0.5, and the slopes were less negative than -1. Importantly, these slopes are complementary to those in **Figure 6**, such that their arithmetic difference equals 1. This means that a slope less than unity in the *δ*_tissue_ vs. *δ*_food_ regression necessarily corresponds to a slope greater than -1 in the Δ vs. *δ*_food_ trend. Both findings indicate partial compensation of dietary SIR variation in tissue, confirming that the effect observed by Caut et al. aligns fundamentally with our slope analysis.

## DISCUSSION

A review of several representative datasets, each involving at least four distinct diets fed to mammals and fishes, revealed a consistent failure of the basic assumption underlying the Δ- based diet-tissue paradigm (2): the requirement for a unitary slope in the linear regression of (*δ*_tissue_, *δ*_food_) was never even approximately met, despite often very high R^2^ values. The non-unitary slopes always result in a Δ-equilibrium point where Δ crosses zero and changes sign — a clear violation of the principle of isotope neutrality, which assumes a quasi-constant Δ that does not change sign.

This inconsistency is not unique to our study. For example, Felicetti et al. reported a slope of 0.42 (R^2^ = 0.88) for carbon in bear plasma, with a Δ-equilibrium at -25.9‰, and a slope of 0.88 (R^2^ = 0.98) for nitrogen (Δ-equilibrium at 6.0‰). Even for sulfur, the slope was 0.78 (R^2^ = 0.98, Δ-equilibrium at 0.9‰). Despite numerous examples of below-unity slopes, the literature offers no convincing explanation for this phenomenon. Perga & Grey suggested that differences in dietary protein content might enhance the negative correlation between Δ and diet [Perga & Grey, 2010], but if this were the case, the nitrogen slope should be similar to or lower than that of carbon; instead, it is much higher.

While it is conceivable that a lower-than-unity slope could arise from data scatter around the fitted line, our Monte Carlo simulations using a noisy *y* = *x* + Const function showed that sub-unity slopes occur about half the time, but so do slopes above unity, resulting in an average slope close to 1. Thus, the persistently observed sub-unity slopes in these studies are real and not artifacts.

A below-unity slope in the *δ*_tissue_ vs *δ*_food_ regression indicates the presence of a compensatory mechanism in tissues that dampens food-induced deviations of SIRs from the Δ-equilibrium. This damping effect causes SIRs in tissues to converge toward the equilibrium value, as vividly illustrated in **Figures 2** and **4**. As a result, the farther a tissue’s SIR is from the Δ-equilibrium, the larger the apparent Δ value becomes: strong enrichment of heavy isotopes leads to a highly negative Δ, while strong depletion yields a large positive Δ.

Empirical evidence supports this pattern. In a study where mice were fed yeast containing 80% ^13^C for extended periods, the ^13^C content in mouse tissues remained substantially below that of the diet, even after 127 and 234 days [Gregg CT et al., 1973]. No organ, including the liver (the most metabolically active), approached the dietary ^13^C level; the highest recorded was 74.6% in liver at day 127, dropping to 62.9% at day 234. Even a fetus conceived during the experiment had only 64.5% ^13^C. These results suggest that tissues with strongly deviating SIRs may gradually drift closer to the Δ-equilibrium over time. The average ^13^C discrimination factor in this study was extremely negative (<-200‰), and the slope of the *δ*_tissue_ vs *δ*_food_ regression was about 0.75, consistent with the slope of approximately 0.6 for carbon observed in rats [Caut S et al., 2008a].

At the opposite extreme — deep ^13^C depletion — Δ-equilibrium should produce large positive Δ values. While there are no rodent studies on such depletion, recent experiments with *C. elegans* worms fed *E. coli* bacterai grown in isotopically depleted media showed significant enrichment in heavy carbon in worm proteins compared to the diet [Zhang X et al., 2024].

Notably, even in the foundational study by DeNiro and Epstein, mouse tissue *δ*^13^C values shifted in opposite directions depending on whether *δ*^13^C food was above or below approximately -21‰ [DeNiro MJ & Epstein S, 1978]. Although the authors did not discuss this obviously contradicting the quasi-constant Δ fact, they cautioned about the intrinsic inaccuracy of their approach and recommended its use only when dietary ^13^C differences were large (e.g., terrestrial vs aquatic diets, or C3 vs C4 plants). It is now apparent that this inaccuracy stems less from individual biochemical variability and more from the deviation from the unity slope implicitly assumed in the traditional diet-tissue paradigm.

Two key questions emerge from the recognition that tissue SIRs converge toward a Δ-equilibrium: the origin of this compensatory phenomenon and its practical application. Regarding the SIRs convergence, its underlying cause remains unresolved and will likely be the subject of future research and debate. While Caut et al. identified the negative correlation between Δ and *δ*_food_, they did not propose an explanation. Analysis of regression slopes (as in **Figure 6**) suggests that tissue SIR-converging processes compensate for roughly half of the deviation of *δ*_food_ from Δ-equilibrium. One plausible explanation is evolutionary adaptation, where biochemical processes have adjusted to average food SIR values over time. Another, less conventional, hypothesis is provided by isotopic resonance theory, which predicts such convergent behavior [Zubarev RA, 2011]. Regardless of the mechanism, these findings challenge the traditional view of isotopes as neutral tracers and instead support the idea that stable isotopes actively influence element assimilation.

The empirical implications of this phenomenon call for a revised paradigm, which can be captured by a linear regression model:

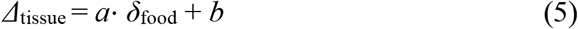

Here, *a* serves as the isotope assimilation factor and *b* is a standard-dependent intercept. As an initial approximation, values for these parameters can be taken from **Figure 6**, but further research is needed to refine them and define their applicability. Alternatively, one can specify the Δ-equilibrium position (e.g., *δ*^13^C = −20.4‰, *δ*^15^N = 11.4‰) and draw a regression line through this point with the empirically determined slope (e.g., *a* = 0.46 for *δ*^13^C, *a* = 0.53 for *δ*^15^N as in **Figure 6**). The intersection of this line with a horizontal line representing a given *δ*_tissue_ value will yield the corresponding *δ*_food_.

This approach provides a statistically robust and mechanistically informed framework for interpreting stable isotope data in diet studies, moving beyond the limitations of the traditional paradigm.

## CONCLUSIONS

Since DeNiro and Epstein first introduced the Δ-based approach (2), it was recognized that Δ is tissue-specific, and later research demonstrated that Δ can also be diet-specific, especially for omnivores. Numerous additional factors — such as trophic position, food chain length, energy flow and sources, and the C/N ratio of the diet — have all been shown to influence Δ [Caut S et al., 2008a]. Despite these accumulating limitations, the diet-tissue paradigm (2) remains dominant in the field, even as practitioners continue to caution that isotope discrimination factors are not yet fully understood (e.g., [Veselý L et al., 2024]). Many published studies begin by affirming the paradigm’s status, only to present new data that challenge its core assumptions.

The central issue is whether the growing body of inconsistencies is now sufficient to warrant a shift to a new paradigm, such as the regression-based model (5). Our analysis underscores a mathematically straightforward but often overlooked point: the assumption of a quasi-constant Δ only holds if the regression slope between *δ*_tissue_ and *δ*_food_ is exactly one. Our review of published datasets shows that this condition is frequently unmet, undermining the universal applicability of the traditional paradigm.

This phenomenon, evident even in the foundational work of DeNiro and Epstein, appears to have been repeatedly observed but rarely highlighted over the past five decades. The notable exception is the attempt by Caut et al., which, while bold, was not fully convincing due to unaddressed spurious correlations. Had similar analyses to those presented here been applied earlier, the paradigm of quasi-constant Δ might have been replaced much sooner by a more realistic model.

It is important to acknowledge that our findings are not universal; cases of regression slopes equal to or greater than unity do exist. For example, our previous research found unexpectedly high deuterium levels in (hydroxy)proline of bone collagen in some marine mammals — an anomaly that cannot be explained by diet alone [Gharibi H et al., 2022]. Nevertheless, the preponderance of well-conducted studies reporting slopes well below unity suggests that tissue SIR convergence is a widespread phenomenon.

In essence, these findings imply that isotopes actively participate in biochemical reactions, influencing both tissue isotopic composition and the enzymes involved, with kinetic feedback mechanisms driving SIR convergence. This feedback fundamentally challenges the notion of isotopes as passive tracers and invalidates the concept of a quasi-constant Δ. The recognition of this active role not only resolves longstanding inconsistencies but also opens exciting new avenues for both theoretical and experimental research in stable isotope science.

The field stands on the threshold of a new era, with the potential for transformative advances in understanding isotope dynamics and their applications in ecological and biochemical studies. The golden years of stable isotopes may yet to come!

## Notes

### Competing Interest Statement

The authors have declared no competing interest.

